# Wnt signaling regulates passive cell competition in the retina by inducing differential cell adhesion

**DOI:** 10.1101/2022.10.27.514088

**Authors:** Xuanyu Min, Yingyu Mao, Hao Wu, Josh Bock, Chenqi Tao, Xin Zhang

**Author notes:** University of Pittsburgh Medical Center (Xuanyu Min).

## Abstract

Self-assortation of progenitor cells during development is essential for establishment of distinct tissue identity. This is exemplified in the eye, where the early optic cup is divided into the neural retina (NR) in the center and the ciliary margin (CM) in the periphery. Previous studies have demonstrated that Wnt signaling is required for specification of the CM, but here we show that genetic ablation of Wnt signaling mediator β-catenin in the peripheral optic cup failed to prevent the formation of the CM-derived ciliary body and iris in adult animals. Mosaic analysis revealed that this was only partially due to loss of adherens junctions among β-catenin deficient cells, which were preferentially excluded from the CM. Even in β-catenin mutant cells that can maintain adherens junctions, their inability to mediate Wnt signaling resulted in a change from P-cadherin to N-cadherin expression. We showed that this cadherin switch was sufficient to segregate otherwise identical cells into separate clusters. As a result, the ciliary body and iris were still formed after inactivation of Wnt signaling in the peripheral retina. These results showed that the dual functions of β-catenin in adherens junction and Wnt signaling are required for the passive cell competition to constitute retinal compartments.

## Introduction

Spontaneous organization guided by intrinsic genetic programs is fundamental to development and functionalities of organs. Within each tissue boundary, this entails self-assortment of relevant cell types as well as repulsion of unwanted intruders. This feat is particularly remarkable considering that the multitude of cell populations may be derived from common progenitors residing at the same anatomical locations during organogenesis. The nature of competitive interactions among progeny cells in establishing their specific tissue niches is an enduring question in biology. A molecular understanding of the underlying mechanism is important for developing stem cell-based regeneration and therapy.

The vertebrate eye is an embodiment of the self-organizing principle in organ development. Millions of neurons and supporting cells that constitute the poster retina can trace their origins to the same progenitors in the optic vesicle, yet they are strictly segregated into adjacent domains of neural retina (NR), ciliary margin (CM) and retinal pigmented epithelium (RPE) (Fig. 1A).

**Figure 1.**
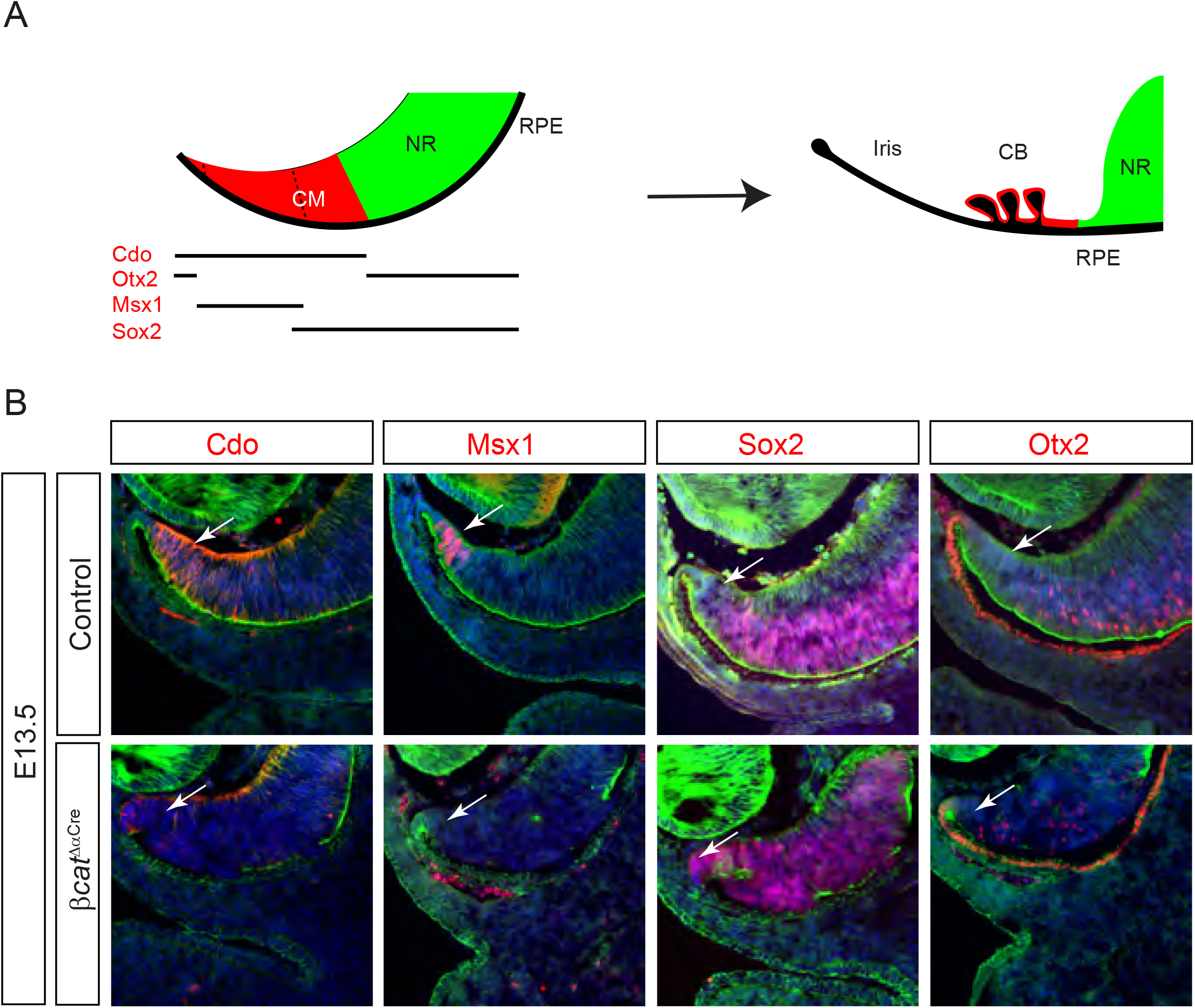
Genetic deletion of β-catenin disrupted the CM development. **(A)** A schematic diagram of CM development. The mouse retina is divided into the central neural retina (NR) and the peripheral ciliary margin (CM), the latter can be further separated into three zones marked by expression of Cdo, Otx2, Msx1 and Sox2. The CM gives rise to the adult iris and ciliary body of the eye. **(B)** Loss of β-catenin in *βcat*^*ΔαCre*^ retinae led to the expansion of Sox2 and Otx2 in the NR at the expense of Cdo and Msx1 expression in the CM region.

Remarkably, such an orderly compartmentalization can be recapitulated entirely in vitro by pluripotent stem cells, which spontaneously differentiate into a broad spectrum of ocular cell types organized into a well-structured retinal organoid. This spatial division occurs despite considerable cell movement found in developing eyes from fish to bird (Casey et al. 2021), in which the CM not only gives rise to the ciliary body and iris, but also serves as a stem cell niche to replenish the NR. Even in mammals, there is evidence that progenitor cells residing in the CM may migrate tangentially into the NR to differentiate into retinal neurons. How these neural and non-neural progenitor cells in the CM eventually home in their respective domains remains an unanswered question.

We have previously shown that the mammalian CM is specified by a binary code of FGF and Wnt signaling, whose relative balance determine the retinal progenitor cell fate in a phase transition like manner. In support of this model, we and others showed that abrogation of Wnt secretion from the lens or Wnt signaling effector β-catenin in the peripheral retina abolished CM development (Liu et al. 2007; Fujimura et al. 2009; Westenskow et al. 2009; Bharti et al. 2012; Balasubramanian et al. 2021). Conversely, constitutive activation of β-catenin in the NR induces the ectopic CM (Liu et al. 2007; Ha et al. 2012). In this study, we followed the consequence of β-catenin deletion in the embryonic retina to the adulthood and unexpectedly found that the CM-derived ciliary body and iris were still formed. In contrast, the NR lost adherens junctions and the basement membrane, leading to impaired astrocytic and vascular networks. This phenotypic disparity between CM and NR compartments correlated with their respective constitutes of wild type and β-catenin deficient cells, which displayed opposite patterns of cadherin expression. By genetically separating adhesive and signaling functions of β-catenin, we showed that canonical Wnt signaling in retinal progenitor cells controlled differential expression of cadherins, which was sufficient to cause cell segregation in vitro. Our results revealed an important role of Wnt signaling in regulating the competitive cell interactions underlying retinal niche development.

## Results

### Genetic ablation of β-catenin disrupted the embryonic CM development but not the postnatal formation of the ciliary body and iris

We have recently showed that the mouse embryonic CM in the peripheral retina can be divided by overlapping expression of molecular markers into the Sox2+/Cdo+ proximal, the Msx1+/Cdo+ medial and Otx2+/Cdo+ distal domains, which resemble the partition of the pars plana and pars plicata of the ciliary body and the iris in the adult eye (Fig. 1A) (Liu et al. 2007; Heavner et al. 2014; Balasubramanian et al. 2021). The CM region can be targeted by *α-Cre* transgene, whose activity is restricted to the peripheral retina as shown by the cistronic GFP reporter. Notably, genetic lineage tracing using an Ai9 Cre reporter showed that *α-Cre* expressing cells eventually extended to a much larger domain in the retina. In *α-Cre; β-catenin*^*flox/flox*^ (*βcat*^*ΔαCre*^) mutants, this led to widespread ablation of β-catenin retinae at E13.5, leading to significant down regulation of both the medial CM marker Msx1 and the pan CM marker Cdo (Fig. 1B, arrows). In contrast, Otx2 and Sox2 expanded from the NR into the CM region. These results confirmed the requirement of β-catenin for embryonic CM development.

Based on the severity of CM defects in E13.5 *βcat*^*ΔαCre*^ embryos, we expected that the CM-derived ocular structures would be absent in adult animals. To our surprise, most newborn *βcat*^*ΔαCre*^ pups exhibited not only normal looking irises (Fig. 2A, arrows), but also double-layered ciliary bodies marked by Otx2 in the pigmented epithelium (PE) and Pax6 and Cdo in the non-pigmented epithelium (NPE) (Fig. 2A, arrows and arrowheads). Similar to controls, these *βcat*^*ΔαCre*^ mutants also expressed water channel protein Aqp1 on basal sides of ciliary bodies and smooth muscle marker SMA in the sphincter pupillae at tips of irises (Fig. 2A, empty arrowheads and asterisks). After the removal of the posterior retina to expose the entire anterior eye segment, most adult *βcat*^*ΔαCre*^ mutants appeared indistinguishable from wild type controls in both their overall structure integrity and the characteristic folding of ciliary bodies (Fig. 2B).

**Figure 2.**
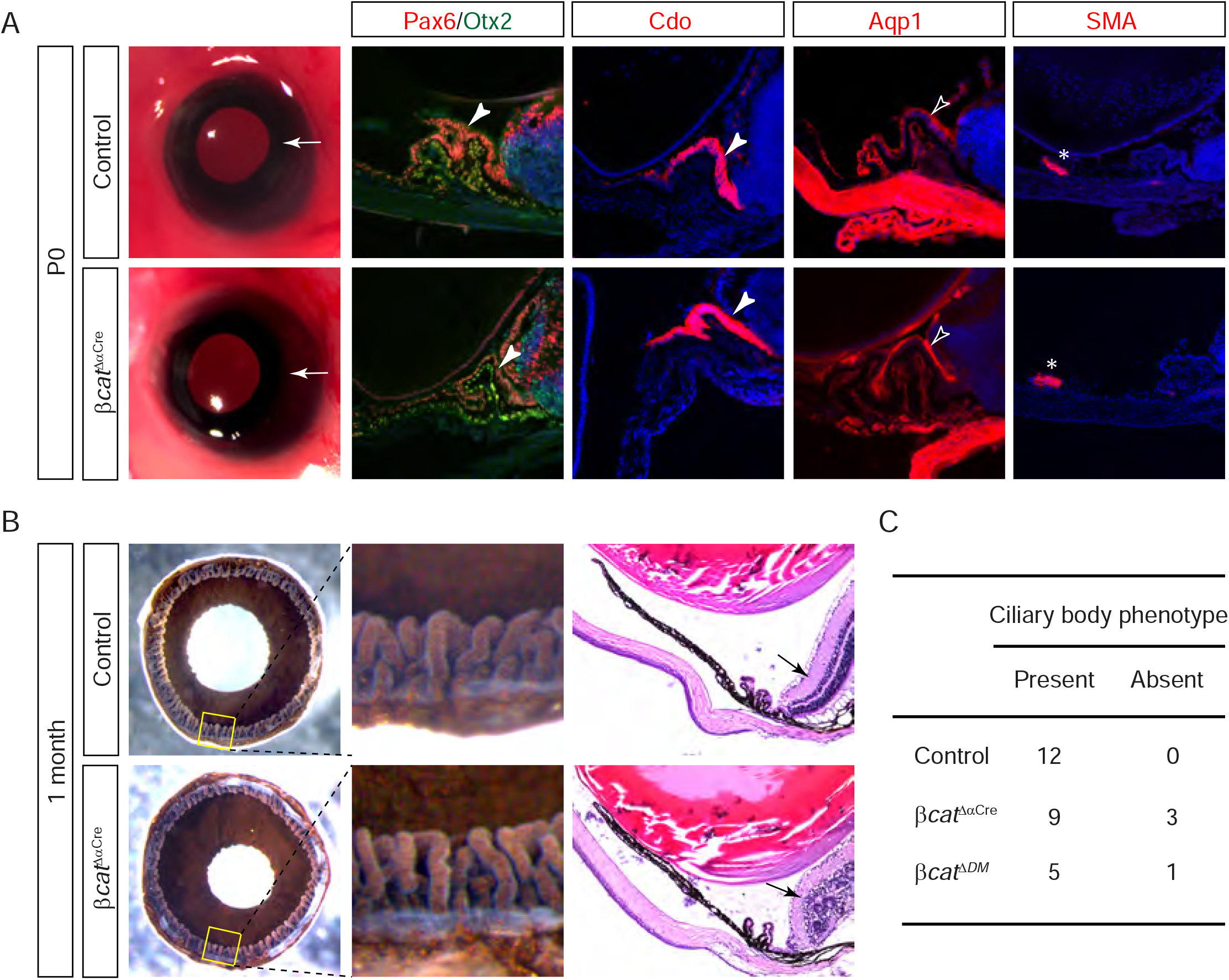
The ciliary body and the iris still formed in adult *βcat*^*ΔαCre*^ mutants. **(A)** Newborn *βcat*^*ΔαCre*^ mutants displayed morphologically intact iris (arrows). The ciliary body specific expression of Pax6, Otx2, Cdo and Aqp1 and the iris specific SMA were also observed (arrowheads). **(B) One** month old *βcat*^*ΔαCre*^ mutants showed well-formed ciliary bodies. **(C)** Statics of the ciliary body (CB) phenotype.

This was further confirmed by histology, which showed that the majority of *βcat*^*ΔαCre*^ mutants retained ciliary bodies and irises (Fig. 2B and C, arrows). Therefore, our embryonic ablation of β-catenin did not abrogate formation of the ciliary body and iris in adult animals.

### Loss of β-catenin resulted in defective retinal basement membrane and impaired networks of astrocytes and endothelial cells

To investigate whether there were any postnatal defects in *βcat*^*ΔαCre*^ mutants, we examined astrocyte migration and angiogenesis, which occur in mouse retina around birth (Tao and Zhang 2014). Specifically, retinal ganglion cells are known to provide the chemotactic signal and migratory tracks for retinal astrocytes, which in turn promote ingression of endothelial cells into the retina (Tao and Zhang 2016). At P4, Brn3a+ retinal ganglion cells were evenly distributed in control retinae, but they appeared to cluster abnormally in the peripheral retinae of *βcat*^*ΔαCre*^ mutants where *α-Cre* is active (Fig. 3A, dotted lines and arrowheads). By examining Fibronectin expression, which was strong in endothelial cells and weak in astrocytes, we observed that this abnormal region has yet to be reached by the retinal vasculature at P4, but it was instead penetrated by hyaloid vessels from the vitreous (Fig. 3A, arrows). In contrast, there was scarcely any astrocytic network in this region. To confirm these phenotypes, we further stained *βcat*^*ΔαCre*^ mutants for Pax2 and Pdgfra, which labelled the nucleus and plasma membrane of astrocytes, respectively. As shown in Fig. 3B (dotted lines), astrocytes specifically avoided the peripheral retina which was covered by IB4+ hyaloid vessels. These findings revealed that β-catenin deficiency disrupted astrocytic and endothelial networks in the postnatal retina.

**Figure 3.**
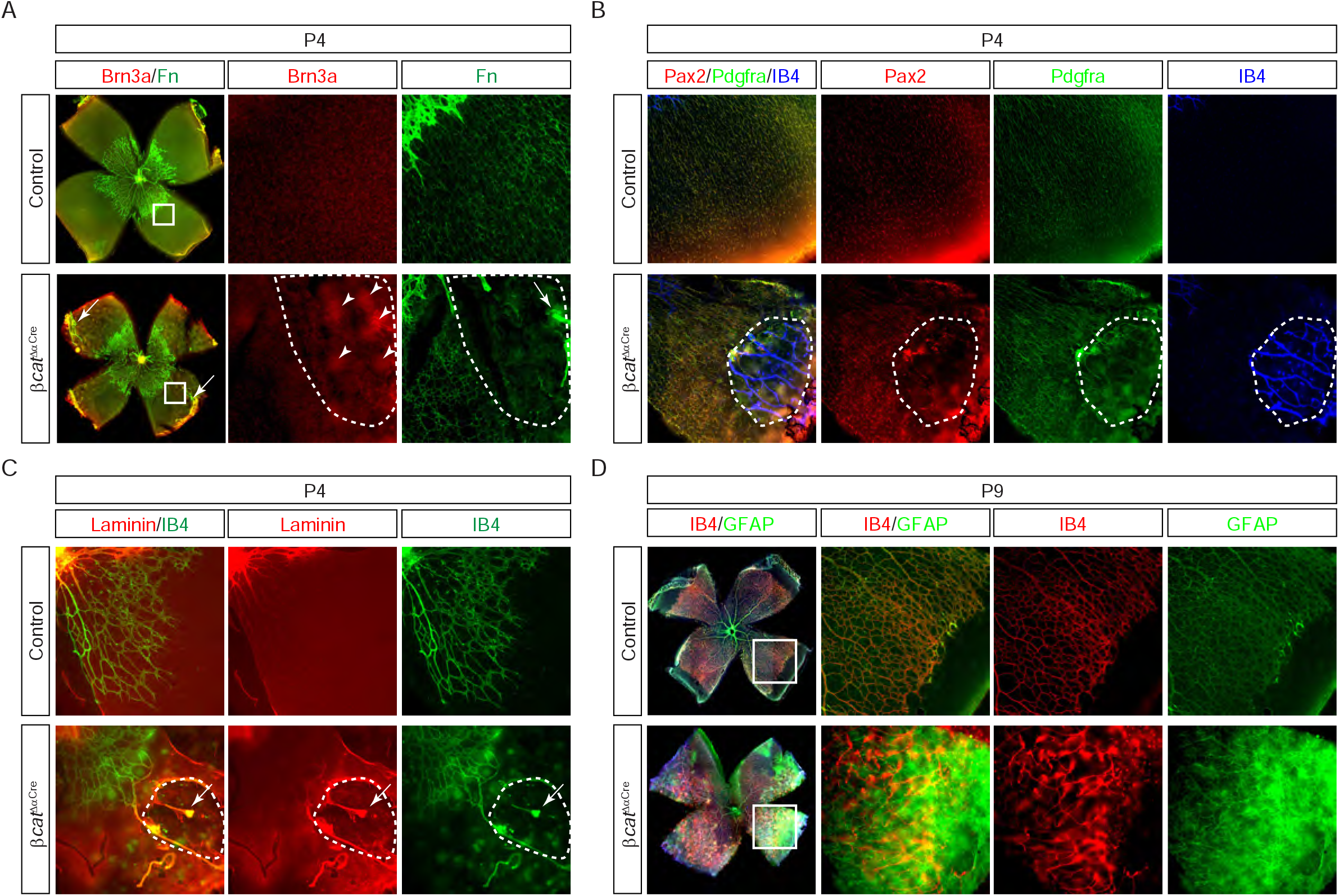
The postnatal development of retinal astrocytes and vasculature was disrupted in *βcat*^*ΔαCre*^ mutants due to the loss of the retinal basement membrane. **(A)** P4 *βcat*^*ΔαCre*^ mutant retinae showed uneven distribution of Brn3a+ retinal ganglion cells (arrows) and disruption of the astrocytic network (dotted lines). These regions were also penetrated by persistent hyaloid vessels (arrow). **(B)** The peripheral retina of *βcat*^*ΔαCre*^ mutants showed sparse astrocytes marked by Pax2 and Pdgfra expression and anormal persistence of hyaloid endothelial cells marked by IB4. **(C)** Laminin staining of *βcat*^*ΔαCre*^ mutants showed extensive breaches in the basement membrane (dotted lines) and associated hyaloid vessels (arrows). **(D)** P9 *βcat*^*ΔαCre*^ mutants showed the irregular vascular network marked by IB4 and retinal gliosis shown by increased GFAP.

Failure in astrocyte migration and overgrowth of hyaloid vessels are indicative of defects in the inner limiting membrane (ILM), the basement membrane of the retina. This is due to the essential roles of the ILM as the migratory substrate for astrocytes and the barrier to prevent infiltration of hyaloid endothelial cells (Tao et al. 2022). Indeed, the ILM appeared as a smooth sheet of Laminin staining in control retinae, but in the peripheral regions of *βcat*^*ΔαCre*^ mutant retinae, it was breached by numerous holes associated with residual hyaloid vessels that were difficult to remove during dissection (Fig. 3C, arrows and dotted lines). At P9 when retinal vessels have finally arrived at the peripheral retina, *βcat*^*ΔαCre*^ mutant retinae displayed extensive irregularities in the vasculature network and significant increase in gliosis marker GFAP (Fig. 3D), both of which have been observed in mouse models with defective ILMs. These results led us to conclude that β-catenin is required for the integrity of the retinal basement membrane.

### β-catenin mutant cells were preferentially excluded from the developing CM

Our study so far has revealed stark phenotypic differences between embryonic and postnatal *βcat*^*ΔαCre*^ mutants. This prompted us to generate a temporal series of β-catenin ablation using the tamoxifen-inducible *Rax-Cre*^*ERT2*^ driver that can target the entire retina (Fig. 4A). Interestingly, induction of Cre activity in *Rax-Cre*^*ERT2*^; *β-catenin*^*flox/flox*^ (*βcat*^*ΔRaxCre*^) at E10.5 only led to modest reduction in β-catenin staining at E14.5. As a result, there was no obvious defect in the localization pattern of either E-cadherin or N-cadherin, except a few breaches at the ventricular side of the posterior retina. Consistent with this, *βcat*^*ΔRaxCre*^ mutants still maintained the typical CM^high^-NR^low^ gradient of Pax6 in the retina and the specific expression of Cdo and Msx1 in the CM. Therefore, the Cre-mediated knockout in the retina led to considerable delay and mosaicism in β-catenin ablation. We next shifted tamoxifen induction one day earlier to E9.5 and harvested the samples at E14.5. Amidst the mosaic loss of retinal β-catenin staining, the remaining β-catenin positive cells retracted their apical plasma membranes, pulling newly differentiated Otx2-expressing neurons away from the retina ventricle (Fig. 4B, arrowheads). This led to formation of evenly spaced cellular rosettes except in the CM region, which remains exclusively occupied by β-catenin positive cells (Fig. 4B, dotted lines). Consequently, the CM specific patterns of Pax6, Cdo and Msx1 expression were still maintained (Fig. 4B, arrows). Considering the ubiquitous nature of the *Rax-Cre*^*ERT2*^ activity in the retina as shown by the Cre reporter (Fig. 4A), we concluded that β-catenin deficient cells in *βcat*^*ΔRaxCre*^ mutants may have been actively excluded from the CM.

**Figure 4.**
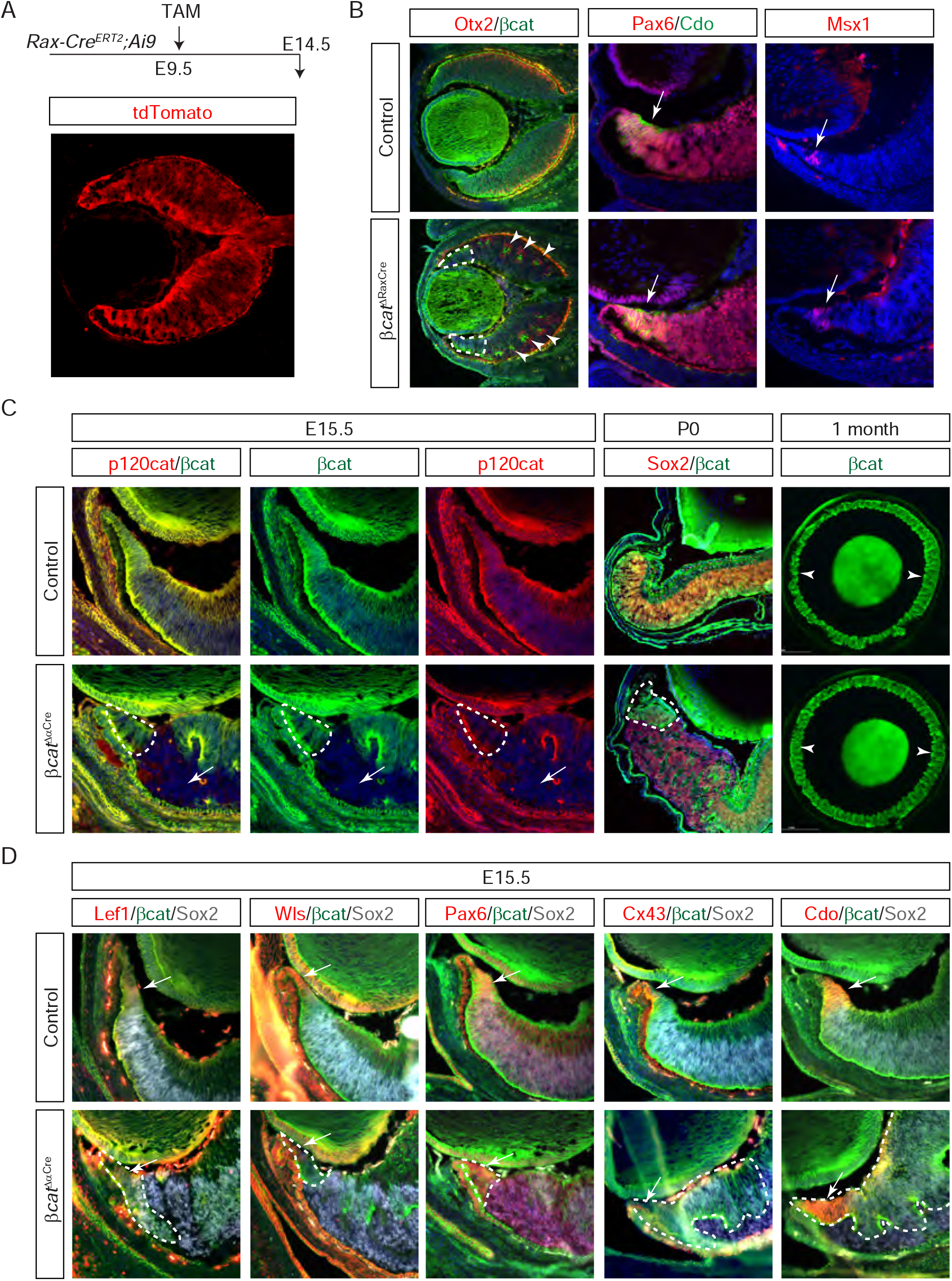
β-catenin deficient cells were excluded from the CM. **(A)** Tamoxifen induction of *Rax-Cre*^*ERT2*^ driver at E9.5 led to widespread activation of *Ai9* tdTomato reporter in the E14.5 eye cup. **(B)** β-catenin deficient cells formed retinal rosettes in *βcat*^*ΔRaxCre*^ mutant retina (arrowheads), but they were excluded from the CM region (dotted lines). As a result, the CM specific expression of Pax6, Cdo and Msx1 were still preserved (arrows). **(C)** In contrast to β-catenin mutant cells that lost p120 catenin staining, wild type cells maintained adherens junction and persisted in the CM from E15.6 to P0 (dotted lines) and in the ciliary body at 1 month (arrowheads). **(D)** Persistent wild type cells in the CM supported active Wnt signaling protein Lef1 and Wls and they expressed the CM specific markers Pax6, Cx43 and Cdo.

We next determined whether this spatially restricted pattern of β-catenin mosaicism was also present in *βcat*^*ΔαCre*^ mutant retinae. Unlike the scattered deletion pattern observed earlier at E13.5 (Fig. 1B), β-catenin was now enriched in the CM at E15.5, where it colocalized with p120 catenin, an important cadherin binding protein required for the stability of adherens junction (Fig. 4C, dotted lines). By contrast, the adjacent NR consisted of random cell aggregates either purely positive or negative for β-catenin and p120 catenin, which explained the formation of cellular rosettes in this region (Fig. 4C, arrows). During the postnatal growth from P0 to one month, the ciliary body region continued to be occupied by β-catenin positive cells, demonstrating the persistence of the β-catenin enrichment (Fig. 4C, dotted lines and arrowheads). As expected from the crucial role of β-catenin in mediating Wnt signaling, the Wnt response proteins Lef1 and Wls remained active in the peripheral retina, enabling the expression of the CM specific markers Pax6, Cx43 and Cdo (Fig. 4D, dotted lines and arrows). These results suggested that the β-catenin dependent reorganization of retinal progenitor cells supported formation of the ciliary body and iris in *βcat*^*ΔαCre*^ mutants.

### The canonical Wnt signaling directed retinal cell segregation

What can account for the apparent sequestration of β-catenin positive cells to the CM region? One plausible reason is that the presence of β-catenin allowed these cells to establish adherens junctions with their wild type neighbors, which concatenated them together to the distal RPE adjacent to the CM. In contrast, β-catenin deficient cells were driven out of the CM niche because they lacked such intercellular anchors. To test this hypothesis, we separated the adhesive function of β-catenin from its role in mediating Wnt signaling using β-catenin^DM^ mutant, which lacks transcriptional activity but still retains the interaction with the cadherin complex (Fig. 5A) (Valenta et al. 2011). By combining *β-catenin*^*DM*^ with the *α-Cre* and the *β-catenin*^*flox*^ allele, it is expected to disable Wnt signaling in the retina without jeopardizing adherens junctions. Indeed, E15.5 *α-Cre; β-catenin*^*flox/DM*^ (*βcat*^*ΔDM*^) mutants did not exhibit any cellular rosettes, but surprisingly, the expression of Wnt response protein Lef1 and Wls were also maintained (Fig. 5B, arrows). As a result, the CM specific pattern of Pax6, Cdo, Otx2, Msx1 and Sox2 expression were still observed in *βcat*^*ΔDM*^ mutants. To corroborate these findings, we also generated *α-Cre; Lrp5*^*flox/flox*^ ; *Lrp6*^*flox/flox*^ (*Lrp5/6*^*ΔαCre*^) mutants to ablate Lrp5 and 6, two obligatory Wnt receptors. In agreement with *βcat*^*ΔDM*^ mutants, even the removal of these Wnt receptors did not prevent the expression of either the Wnt response protein or the CM specific markers in E15.5 *Lrp5/6*^*ΔαCre*^ retinae. These results suggest that localization of wild type cells to the CM can not be explained solely by the adhesive function of β-catenin.

**Figure 5.**
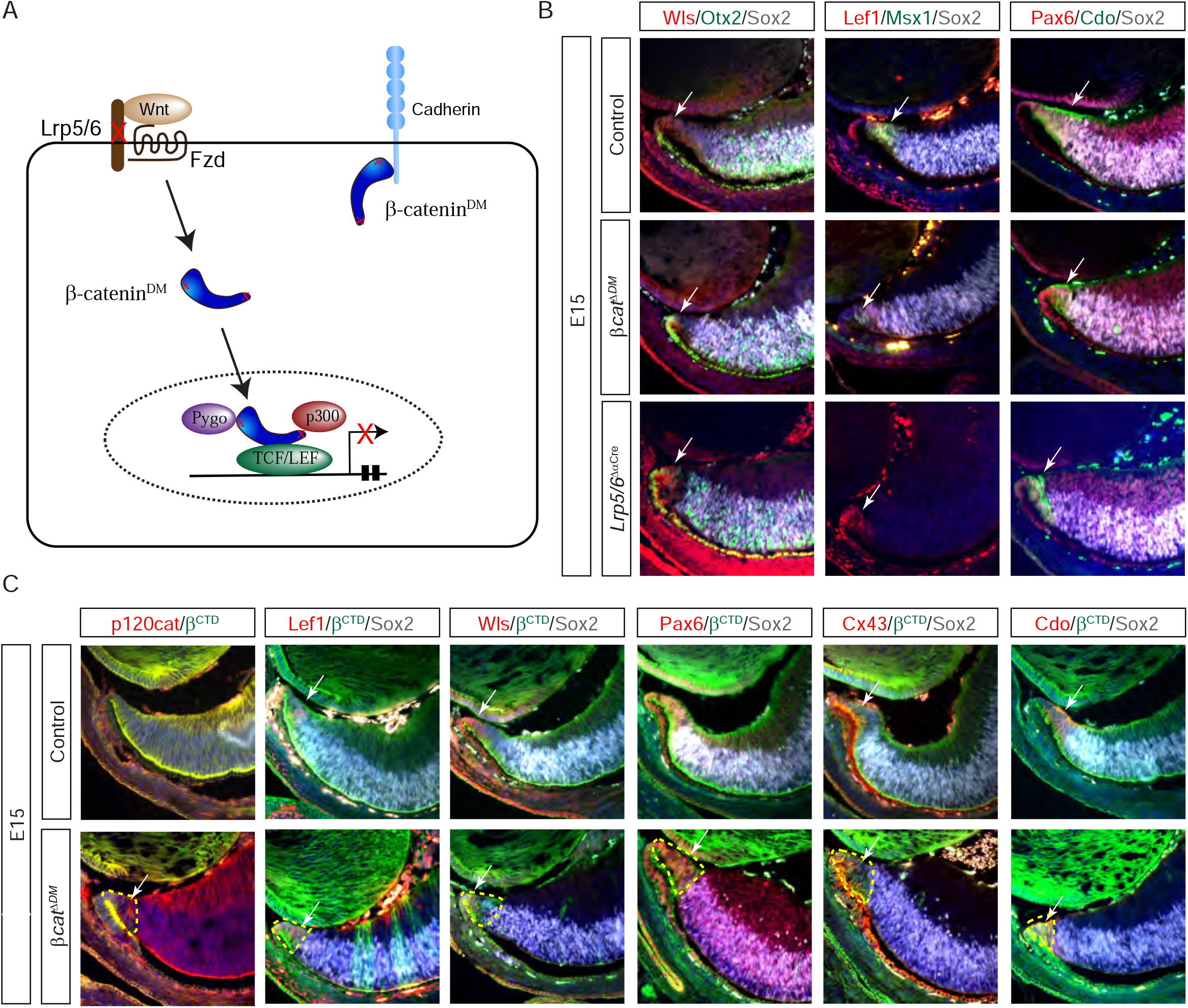
The retention of wild type cells in the CM was independent of the adhesive function of β-catenin. **(A)** A schematic diagram of the dual functions of β-catenin in adherens junction and Wnt signaling. The point mutations in β-catenin^DM^ mutant abolished its transcriptional activity without interfering with its interaction with cadherins. **(B)** Neither inactivation of the β-catenin transcriptional activity in *βcat*^*ΔDM*^ nor ablation of Wnt receptors in *Lrp5/6*^*ΔαCre*^ mutants abolished Wnt signaling proteins Wls and Lef1 or the CM markers Msx1, Pax6 and Cdo (arrows). **(C)** Wild type cells remained enriched in the CM region in *βcat*^*ΔDM*^ mutants, where they supported expression of adherens marker p120 catenin, Wnt signaling protein Lef1 and Wls and the CM specific markers Pax6, Cx43 and Cdo. Wild type cells persisted in the CM from E15.6 to P0 (dotted lines) and in the ciliary body at one month (arrowheads). **(D)** Persistent wild type cells in the CM supported active Wnt signaling protein Lef1 and Wls and they expressed the CM specific markers Pax6, Cx43 and Cdo (arrows and dotted lines).

To resolve why Wnt signaling remained active in *βcat*^*ΔDM*^ mutants, we visualized the mosaic pattern of genetic ablation in E15.5 retinae using an antibody against the C-terminal domain of β-catenin (β^CTD^), which was truncated in β-catenin^DM^ protein. Interestingly, we observed that wild type cells were again enriched in the CM region at the expense of *β-catenin*^*DM/-*^ mutant cells, despite the normal pattern of p120 catenin staining indicative of intact adherens junctions (Fig. 4C). Consistent with the inability of β-catenin^DM^ mutant to transmit Wnt signaling, Wnt signaling markers Lef1 and Wls were only detected within islands of wild type cells, which also supported the expression of the CM markers Pax6, Cx43 and Cdo (Fig. 4C). Therefore, the active Wnt signaling in *βcat*^*ΔDM*^ mutants was the manifest of skewed contribution of wild type cells to the CM region. It further suggests that the canonical Wnt signaling regulates the cellular competition for this retinal niche.

### Wnt signaling induced distinctive pattern of cadherins in the retina

To understand how Wnt signaling may regulate the competitive cellular interactions in the CM, we first considered the cell competition model whereby wild type cells gain growth advantage by hyperproliferation while inducing apoptosis in their less fit mutant neighbors.

However, both the TUNEL assay and cleaved Caspase-3 staining failed to detect any abnormal apoptosis in *β-catenin*^*DM/-*^ mutant cells. On the other hand, the proliferation rate as determined by Ki67 staining was also similar between wild type and mutant cells. In fact, there were overall more mutant cells than wild type cells in *βcat*^*ΔDM*^ mutants, suggesting that Wnt signaling was unlikely to have affected cell fitness within the retina.

While the classic cell competition model emphasizes active elimination of competitors by more fit cells, we next considered the possibility that retinal progenitor cells may engage in a more passive contest for the CM niche. In this alternative model, mutant cells may have been excluded from the tissue niche if they lacked the requisite adhesion to their wild type neighbors due to the change in their surface protein expression. The likely candidates for mediating such differential cell adhesion are cadherins, whose family members may engage in either homophilic or heterophilic interactions (Katsamba et al. 2009; Fagotto 2014). We found that there were three cadherins expressed in the E15.5 retina: R-cadherin was expressed in both the NR and CM, P-cadherin was only detected in the CM region, whereas N-cadherin was predominantly in the NR (Fig. 6A, arrows). Importantly, *β-catenin*^*DM/-*^ mutant cells exclusively expressed N-cadherin, but not P-cadherin, while R-cadherin was expressed at the same level in both wild type and mutant regions (Fig. 6A, dotted lines). The differential expression of cadherins persisted in P3 retinae, where P-cadherin expression was confined to wild type cells in the ciliary body and N-cadherin expression was restricted to *β-catenin*^*DM/-*^ mutant cells in the NR (Fig. 6B, arrows and dotted lines). Therefore, Wnt signaling regulated the distinct expression of P- and N-cadherins in the CM versus the NR.

**Figure 6.**
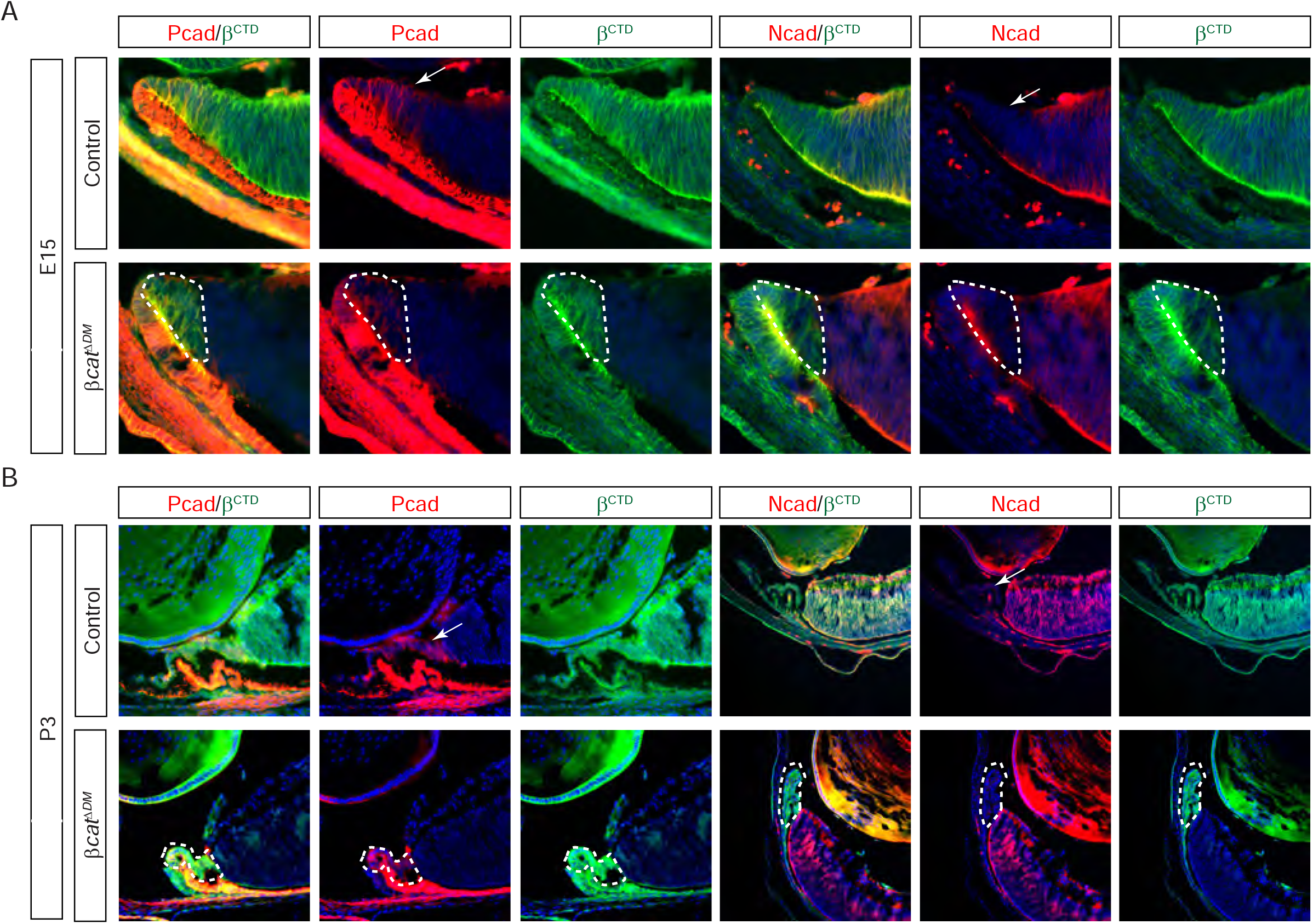
Wnt signaling induced differential expression of cadherins in the retina. **(A)** P-cadherin expression was restricted to the CM region (arrows) where it colocalized with the residual wild type clones (dotted lines) in *βcat*^*ΔDM*^ mutants. In contrast, N-cadherin was predominantly expressed in the NR, which was unaffected in *β-catenin*^*DM/-*^ mutant cells. **(B)** In P3 retina, P-cadherin (arrow) was only detected in the ciliary body which consisted of β-catenin positive cells (dotted lines), while N-cadherin expression was present in the β-catenin deficient NR.

### Differential expression of cadherins promoted passive cell competition to support CM development

Is differential expression of P- and N-cadherin sufficient to induce cell segregation? We first evaluated this possibility based on the biophysical properties of these two molecules. According to the differential adhesion hypothesis (DAH), the aggregation pattern of cells expressing two different cadherins depends on the value of Φ = [*W*(*I, I*)+*W*(*J, J*)]/2-*W*(*I, J*)], where *W* is the work required to separate two cadherin-expressing cells, while I and J denote the type of cadherins. For Φ > 0, the DAH model predicts that I and J cells will form separate homotypic clusters, which are unable to make close contact to each other if *W*(*I, J*) ≈ 0. Previous studies have shown that P-cadherin and N-cadherin have the lowest homophilic dimerization *K*_*D*_s among the class I cadherins, but their heterophilic dimerization *K*_*D*_ was among the highest. As a result, both *W*(*P, P*) and *W*(*N, N*) are far larger than *W*(*P, N*), which is close to zero, rendering Φ > 0. This theoretical consideration suggested that P-cadherin and N-cadherin-expressing cells would spontaneously form separate aggregates without contact.

We next tested the predication from the DAH model using K562 cells, which lacked endogenous expression of cadherins. This allowed us to electroporate them with either P- or N-cadherin constructs tagged with tdTomato or eGFP at their C-terminal ends, respectively. An equal number of P- or N-cadherin expressing cells were mixed by gentle shaking and allowed to aggregate in suspension culture (Fig. 7A). Remarkably, from an initial random distribution of single cells, there emerged aggregates of pure tdTomato and eGFP positive cells with little contact between them (Fig. 7B). This was quantified by measuring the maximum size of aggregates, which showed that homotypic clumps could reach a much larger size than heterotypic ones. We further tested the DAH model by manipulating the strength of adherens junctions. First, we took advantage of the hyperactive Abl signaling in K562 cells, which has been shown to promote adherens junction. We reasoned that inhibition of Abl kinase would reduce the work (*W*) needed to separate two cells in general, leading to a still positive but lower Φ value. Indeed, addition of Abl inhibitor imatinib did not abrogate the homotypic aggregates, but their sizes were smaller. In the second approach, we co-transfected P- and N-cadherin with E-cadherin, which mimicked the in vivo situation where the unique expression pattern of P- and N-cadherin was overlaid with the ubiquitous expression of R-cadherin in the retina. This was expected to strengthen both the homophilic and heterophilic interaction between P/E- and N/E-cadherin expressing cells, so the value of Φ was unchanged. However, as the *W*(*P/E, N/E*) was now increased, the DAH model predicted that these cells should form homotypic aggregates fused together. Indeed, P- and N-cadherin expressing cells still formed pure aggregates, but these cell clumps were now joined to each other, resembling the pattern of adjacent CM and NR in the retina. These results suggested that the combined expression of specific and common cadherins were conducive to the spatial patterning of the retina.

**Figure 7.**
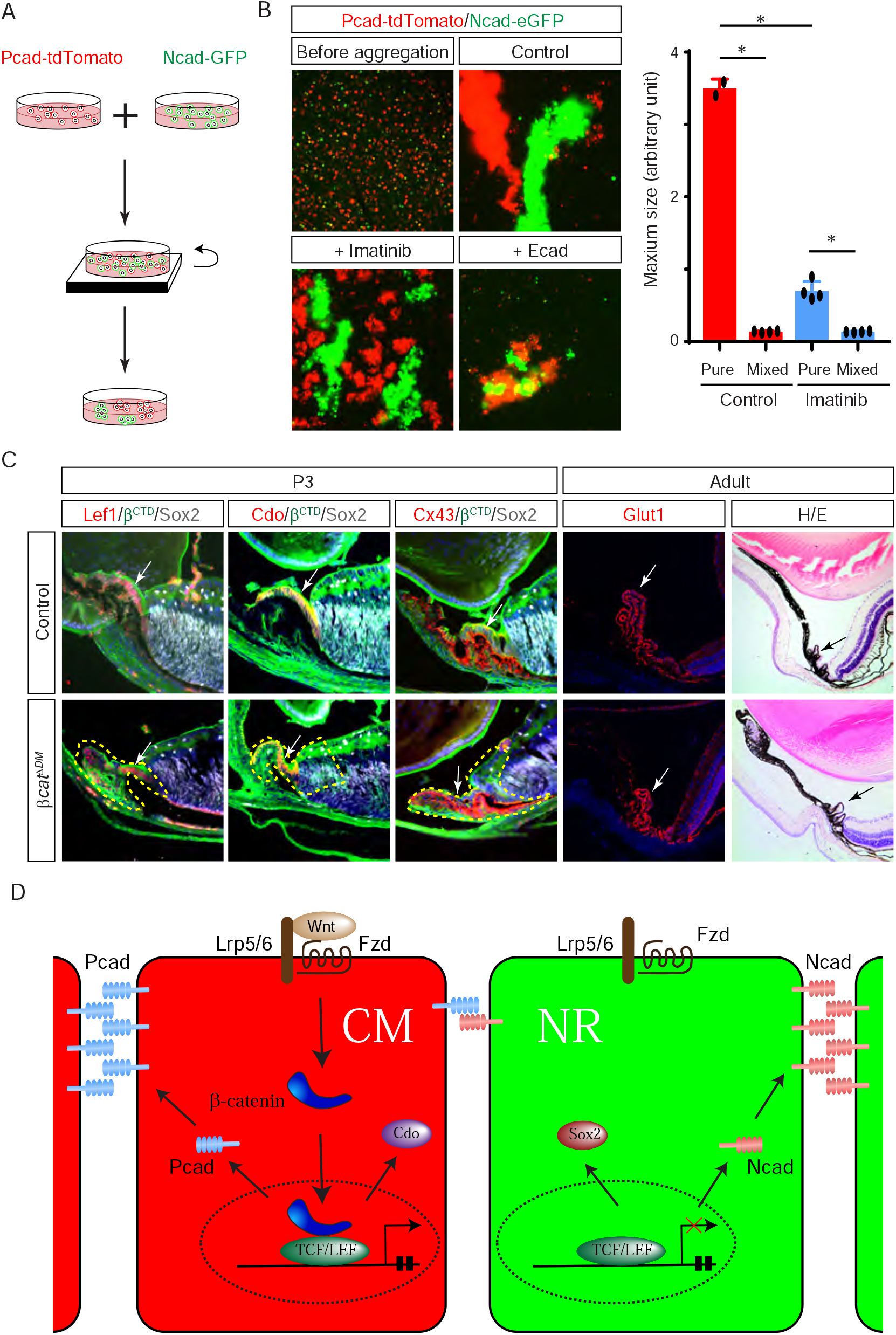
Differential expression of cadherins promoted cell segregation and development of the ciliary body and iris. **(A)** Schematic diagram of the cell aggregation assay. **(B)** An equal mixture of P-cadherin cells marked by tdTomato and N-cadherin cells marked by eGFP spontaneously form pure homotypic aggregates without contact after shaking for 3 hrs. Addition of Abl inhibitor Imatinib reduced the size of the aggregates, while co-transfection of E-cadherin led to embedding of these cell clumps. *One way ANOVA, P<0.001. **(C)** The ciliary body in *βcat*^*ΔDM*^ mutants were occupied by β-catenin positive cells at P3, which expressed Lef1, Cdo and Cx43 (arrows and dotted lines). This led to the formation of Glut1 positive ciliary body in adults (arrows). **(D)** The model of differential cell adhesion directed by Wnt signaling. Wnt signaling induces expression of P-cadherin in the CM cells, leaving the NR cells to express N-cadherin. Because the homophilic affinity of P- and N-cadherin far exceeds their heterophilic affinity, the P- and N-cadherin positive cells spontaneously segregate into the CM and NR regions, respectively.

Lastly, we examined whether the differential cell segregation directed by Wnt signaling cells can support formation of the ciliary body and the iris. In P3 *β-catenin*^*DM/-*^ mutants, wild type cells labeled by β^CTD^ antibody expressed Lef1, Cdo and Cx43 in the ciliary body as expected (Fig. 7C). In adults, *β-catenin*^*DM/-*^ mutants also maintained the stereotypical morphology of the ciliary body and the specific expression of Glut1. Similarly, P3 *Lrp5/6*^*ΔαCre*^ mutants exhibited the expression of Vsx2, Cx43, Glut1, Cyclin D3, Otx2 and Pax6 in the ciliary body and SMA and Cav3 in the iris. The ciliary body and iris in adult animals were further confirmed by expression of Pax6, SMA and Cav3 and by histology. These results showed that the CM specific enrichment of wild type cells directed by Wnt signaling mutants were able to form these peripheral ocular structures.

## Discussion

In this study, we investigated the paradoxical observation that genetic ablation of β-catenin in the retina disrupted the embryonic development of the CM but not the postnatal formation of the ciliary body and iris. Despite the widespread Cre activity, we observed that the mosaic deletion of β-catenin coalesced into separate wild type aggregates in the CM and mutant clones in the NR. While β-catenin deficiency in the NR led to dysregulation of astrocyte migration and angiogenesis, wild type cells were able to respond to Wnt signaling in the CM to form the ciliary body and iris. This spontaneous segregation of wild type and mutant cells were only partially promoted by the loss of adhesive interaction in β-catenin mutant cells, as inactivation of Wnt signaling alone can also lead to enrichment of wild type cells in the CM. Mechanistically, we found Wnt signaling induced P-cadherin and suppressed N-cadherin in the retina. Due to their preference to homophilic rather than heterophilic interaction, these two cadherins were sufficient to drive spontaneous cell segregation in a cell-based assay. Thus, differential cell adhesion directed by Wnt signaling provides a mechanism for the competitive cellular interaction in establishing tissue niche during development.

The classic cell competition is originally discovered in studies of mosaic clones in model systems, where more fit cells eliminate their less fit neighbors by causing their death. It has since been found to underlie myriad biological processes, including organ development, tissue homeostasis and cancer growth. However, it has become increasingly clear that cells can engage in many modes of competitive interactions. On one hand, these interactions may be classified based on direct contest involving cell-cell contact versus scrambling for factors in the microenvironment. On the other hand, they may be divided into costly action involving energy expenditure versus inexpensive action based on cell intrinsic property. The combination of these orthogonal modes can explain many competitive cellular interactions in nature, but interestingly, it is noted that a clear example of the so called “inexpensive contest” has not been observed. Our work helps to fill this gap by showing that the differential cell adhesion in the CM fits such criteria: It is a direct contest based on cell-cell contact, but it is also a passive cellular interaction without causing cell death. These findings thus break new ground on the mechanism of cell competition in organ development.

The differential adhesion hypothesis (DAH) is an elegant biophysical model to explain the spontaneous cell sorting and tissue layering in developing embryos. It posits that the strengths of intercellular interactions determine cell aggregation and cell sorting. By measuring adhesion and surface tension at the single-cell level, the model has found ample success in predicting the behavior of cell aggregation in vitro. However, much less is known how these properties affect cell sorting in vivo. Here we showed that the dimerization affinities of P- and N-cadherin obtained from measuring recombinant protein were sufficient to account for the aggregation pattern of cells expressing these two cadherins. Importantly, our results showed that the DAH prediction also fit closely the mosaic pattern of P- and N-cadherin expressing cells in the developing retina. By manipulating the strengths of adherens junctions using Abl inhibitor or expressing another cadherin, we showed that the more complex pattern of cell aggregation can be generated. Of particular interest is that a combination of unique and common cadherins were able to generate homotypic aggregates fused together, which more closely resemble in the tissue compartmentation in retina. Our result may spur further investigation into the combinatorial actions of adhesion molecules to assemble complex tissue patterns.

Another element of interest concerns the indirect role of β-catenin in the NR on angiogenesis and astrocyte migration in the retina. We and other have previously shown that the basement membrane of the retina, also known as the inner limiting membrane (ILM), serves as the migratory substrate for astrocytes, which secreted angiogenic factors to recruit endothelial cells. In β-catenin deficient retina, we observed severe disruption in the integrity of the ILM, which explained the defects in astrocyte migration and angiogenesis. This result suggested that β-catenin may be required for the assembly and/or integrity of the basement membrane. As a key component in adherens junction, β-catenin links classic cadherins to actin and microtubules, supporting epithelial tissue integrity and maintaining the apical-basal polarity (Harris and Tepass 2010). Its loss in the retina destabilizes cell-cell adhesion, altering cell orientation and leading to rosettes formation. The polarity defect in the retina may have led to dysregulation of the intracellular transport of basement membrane materials or mis localization of the extracellular matrix receptors, both are required for the assembly and maintenance of the basement membrane. Previously studies have demonstrated that integrin signaling plays an important role in establishing adherens junction. Our results in the retina suggest that that the cell-cell adhesion can also have a strong impact on the cell-matrix adhesion.

In summary, we have identified that the unexpected normal ciliary body and iris in β-catenin mutants were generated by a passive cell competition mechanism that may have broader implications for tissue layering and compartmentation. Our study showed that extracellular signaling can induce differential adhesion to direct cell movement, which facilitates appropriate progenitor cells to constitute specific tissue niches. This mechanism not only sheds light on spontaneous cell segregation in organ development, but it may also be harnessed to support the stem cell-based tissue regeneration.

## Methods and Materials

### Mice

All animal procedures were performed according to the protocols approved by the Columbia University’s Institutional Animal Care and Use Committee. *β-catenin*^*fl3*^ was obtained from Dr. Stavroula Kousteni (Columbia University), *Pax6 α-Cre (α-Cre)* from Dr. Nadean Brown (Children’s Hospital Research Foundation), and *Ai9, β-Catenin* ^*flox*^ from Jackson lab (Ashery-Padan et al. 2000). Mice were maintained on a mixed genetic background. At least three animals were analyzed for each of the crosses described. We did not observe phenotypic variations between Cre-heterozygous controls and no-Cre homozygous controls. These two genotypes are hence described together as controls. Tamoxifen was prepared in corn oil at 10 mg/mL and kept at 4°C. Intraperitoneal injections were performed at 100 mg/g of mouse body weight. Three consecutive injections were carried out to achieve maximum Cre induction. Mouse embryos were harvested at different developmental stages from timed pregnant females, followed by genotype and 4% paraformaldehyde (w/v) fixation.

### Immunohistochemistry

Histology and immunohistochemistry were performed on the paraffin and cryosections as previously described (Carbe and Zhang 2011; Carbe et al. 2012). Whole retina fixed in 4% PFA or 10 μm rehydrated cryosections were blocked with 10% normal goat serum (NGS) for 1 hour at room temperature and incubated with primary antibody overnight at 4°C. After washing with PBS, samples were incubated with 2^nd^ fluorescent-conjugated antibody in 2% BSA for 1 hour at room temperature in dark. Isolectin GS-IB_4_ (IB4) conjugated with Alexa Fluor 488 (#I21411, Thermo Fisher Scientific) was applied to visualize the vasculature. Samples were washed and mounted with n-propyl gallate (NPG) anti-fading reagent and examined under a Leica DM5000-B fluorescent microscope. Antibodies used were: anti-GFAP (#Z0334, 1:1000, Dako), anti-Laminin (#L9393, 1:1000, Sigma-Aldrich), anti-Pax2 (PRB-276P, 1:200, Covance), anti-Pdgfra (#558774, 1:1000, BD Pharmingen) and anti-SMA (M085129-2, 1:1000, Agilent). All commercial antibodies are validated by vendors. At least three embryos of each genotype were stained for each marker.

### K562 cell aggregation assay

3 × 10^5^ cells in 20 µL transfection buffer were mixed with 1µg plasmids and electroporated with a 4D nucleofector (Lonza Bioscience) following the preset protocol for K562. >90% transfection efficiency was achieved after 24 hours of incubation. 24 hours after electroporation, 200 µL of 500 µL cells expressing either N- or P-cadherin were mixed into one well of a 24-well plate with/without imatinib treatment (10 µM) and then incubated on a rocker at 37ºC 5% CO2 for 3hr. Cells were imaged immediately after incubation.

## Acknowledgements

The authors thank Dr. Stavroula Kousteni for mice and Dr. Sha Zha for help. We thank members of Zhang lab for discussion. The work was supported by grants from NIH (EY017061, EY018868, EY025933 and EY031210 to X.Z.). The Columbia Ophthalmology Core Facility are supported by NIH Core grant 5P30EY019007 and unrestricted funds from Research to Prevent Blindness (RPB). C.T. is a recipient of Jonas Scholar award.

